# Na_v_1.5 single-molecule localization reveals different reorganization modes at cardiomyocyte domains

**DOI:** 10.1101/674275

**Authors:** Sarah H. Vermij, Jean-Sébastien Rougier, Esperanza Agulló-Pascual, Eli Rothenberg, Mario Delmar, Hugues Abriel

## Abstract

Mutations in the gene encoding the sodium channel Na_v_1.5 cause various cardiac arrhythmias. This variety may arise from different determinants of Na_v_1.5 expression between cardiomyocyte domains. At the lateral membrane and T-tubules, Na_v_1.5 localization and function remain insufficiently characterized. We used novel single-molecule localization microscopy (SMLM) and modeling to define nanoscale features of Na_v_1.5 localization and distribution at the lateral membrane, groove, and T-tubules in wild-type, dystrophin-deficient (*mdx*) mice, and mice expressing C-terminally truncated Na_v_1.5 (ΔSIV). We show that Na_v_1.5 organizes as distinct clusters in the groove and T-tubules which density and distribution partially depend on SIV and dystrophin. We found that overall reduction in Na_v_1.5 expression in *mdx* and ΔSIV cells results in a non-uniform redistribution with Na_v_1.5 being specifically reduced at the groove of ΔSIV and increased in T-tubules of *mdx* cardiomyocytes. Na_v_1.5 mutations may therefore site-specifically affect Na_v_1.5 localization and distribution depending on site-specific interacting proteins.

## INTRODUCTION

Proper function of the heart heavily relies on the cardiac sodium channel Na_v_1.5. Na_v_1.5 generates the rapid upstroke of the cardiac action potential; thus, it is pivotal for cardiac excitability, conduction, and muscle contraction^1,2^. Mutations in its gene *SCN5A* have been associated with many cardiac arrhythmias, such as long-QT syndrome type 3, Brugada syndrome, and sick-sinus syndrome^3^. Mutations can cause loss or gain of function of Na_v_1.5, but the wide diversity in cardiac arrhythmias remains unexplained. The key may lie in the distribution of Na_v_1.5 over specific membrane domains – “pools” – of the cardiomyocyte. Na_v_1.5 pools are regulated by different proteins, so a given Na_v_1.5 mutation may affect a specific pool if it disturbs the interaction with a specific protein. Na_v_1.5 expression at the intercalated disc^4^ and at the lateral membrane^5^ is well established (**Figure 1**) while a third pool at the T-tubules is still disputed^6^. At the intercalated disc, where two cardiomyocytes are tightly coupled electrically and mechanically, Na_v_1.5 interacts with ankyrin-G, connexin-43, and N-cadherin, among others^4^. At the lateral membrane, where costameres connect the cytoskeleton to the extracellular matrix (**Figure 1a,b**), Na_v_1.5 interacts with the syntrophin/dystrophin complex, which is important for Na_v_1.5 stability, and CASK, among others^7–9^. The lateral membrane is shaped distinctly with valley-like grooves and hilly crests^10^, and Na_v_1.5 is expressed at both locations in murine cardiomyocytes (**Figure 1a**)^11,12^. T-tubules are deep invaginations of the lateral membrane that originate in the groove. They mainly provide a large surface for calcium handling, and are crucial for the uniform contraction of ventricular cardiomyocytes^13^. Consequently, the majority of the voltage-gated calcium channels Ca_v_1.2 is expressed at the T-tubules^14,15^. The claim of a T-tubular Na_v_1.5 pool has so far only been supported by inconclusive results. By comparing whole-cell *I*_Na_ densities in detubulated and control cardiomyocytes, T-tubular sodium channels were determined to conduct 30% of total sodium current^16^. This is probably a considerable overestimation as this method relies on the membrane capacitance as a measure for cell membrane area, but underestimates the surface area of cardiomyocytes by about 50%^17^. Therefore, the presence of a T-tubular Na_v_1.5 pool remains undetermined.

**Figure 1.**
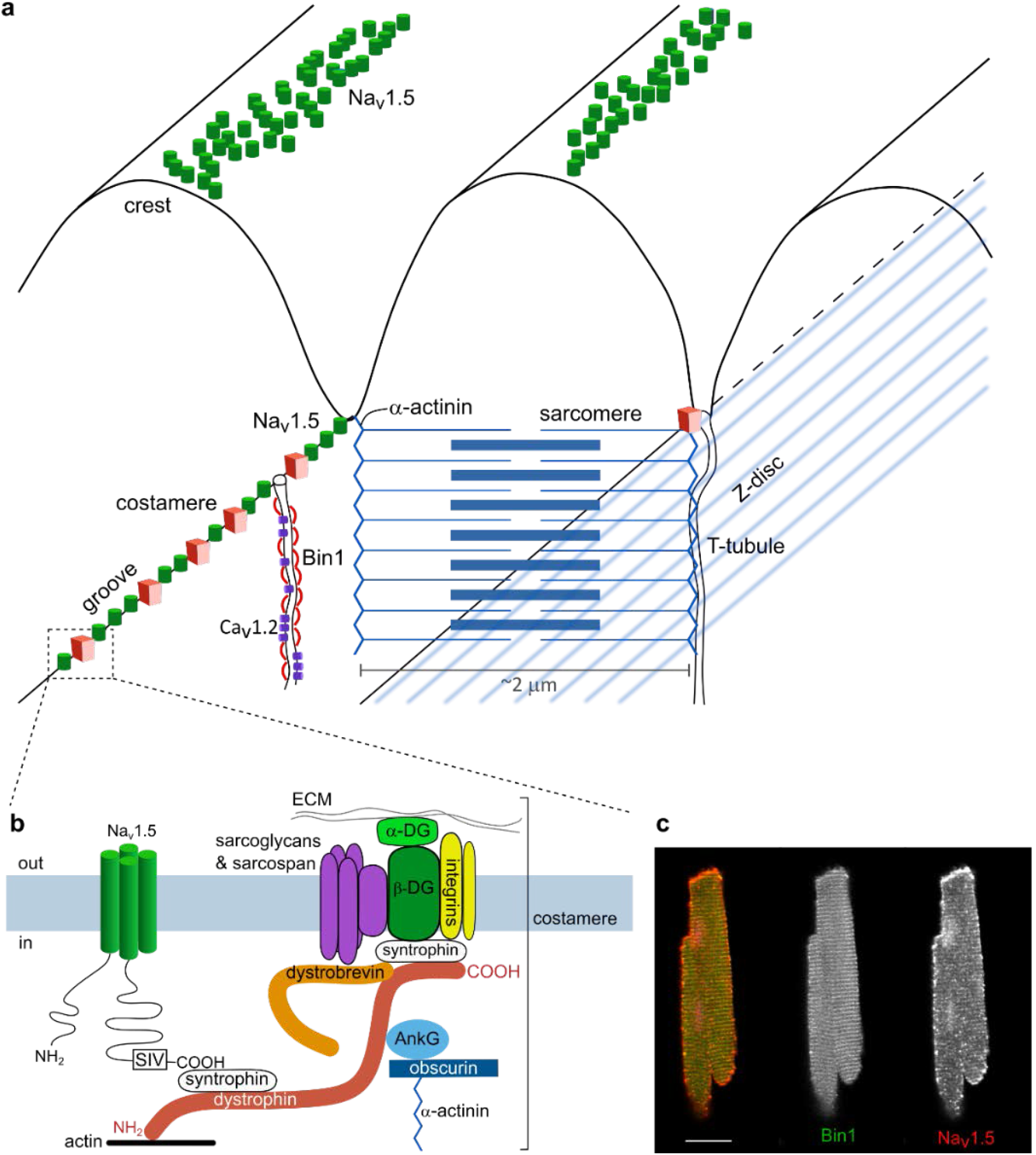
Pools of Na_v_1.5 in the cardiomyocyte. **a**, The lateral membrane has a characteristic profile: at the groove, the membrane is anchored to the sarcomeric Z-line (α-actinin; blue) through costameres (red boxes). T-tubules follow Z-lines and open to the groove. Their membrane is shaped by Bin1 (red curves) and contain, among many other proteins, voltage-gated calcium channels (Ca_v_1.2; purple). The crest has a dome shape. Na_v_1.5 is expressed at the crest and the groove. The amount of Na_v_1.5 in this image serves illustrative purposes only and does not reflect experimental values. **b**, At the groove, Na_v_1.5 (green; left) is associated with the costamere (right). The C-terminal SIV motif of Na_v_1.5 binds syntrophin (white), which binds dystrophin (red). Dystrophin also binds syntrophin associated with the costamere, which contains multiple transmembrane proteins, including integrins (yellow), sarcoglycans (purple), and β-dystroglycan (green). β-dystroglycan connects the sarcolemma with the extracellular matrix (ECM) *via* α-dystroglycan. Dystrophin also links the costamere to the cytoskeleton through its association with actin at its N-terminus and with α-actinin via ankyrin-G and obscurin. **c**, Confocal image of murine ventricular cardiomyocyte stained for Na_v_1.5 (red) and T-tubular marker Bin1 (green), showing Na_v_1.5 expression at the intercalated disc, lateral membrane, and an intracellular punctate pattern coinciding with Bin1 signal. Scale bar 20 μm.

Our current state of knowledge regarding the localization and distribution of Na_v_1.5 at the lateral membrane and T-tubules is based on standard microscopy methods that are limited in resolution and sensitivity. At the T-tubules, previously published immunofluorescence data on Na_v_1.5 show an intracellular striated pattern for Na_v_1.5, but without a T-tubular marker, this pattern cannot be attributed to the T-tubules^18,19^. The respective confocal microscopy techniques moreover lacked resolution to resolve the narrow T-tubular structure^18,19^. At the lateral membrane, cell-attached and scanning ion conductance microscopy (SICM) recordings of the sodium current have provided information on the number of functional channels in a given area at the lateral membrane or specifically at the crest and groove^11,12,20,21^, but they did not provide the molecular features needed to characterize Na_v_1.5 organization at these domains. Moreover, previous studies did not correlate the signal of Na_v_1.5 with other structural markers such as α-actinin for the groove and Bin1 for the T-tubules^18,19,22^. As such, current models of the roles of Na_v_1.5 at the lateral membrane and T-tubules do not represent the true molecular nature of Nav1.5 expression and organization at these domains. To address these issues, we utilize multi-color single-molecule localization microscopy (SMLM)^23^, a super-resolution imaging method, providing the features of Na_v_1.5 organization and related markers with nanoscale resolution. Using this approach, we addressed Na_v_1.5 expression and cluster organization at the T-tubules using the T-tubular marker Bin1^24^, and Na_v_1.5 affinity for the groove using the groove marker α-actinin in isolated cardiomyocytes. We also assessed T-tubular *I*_Na_ by comparing whole-cell *I*_Na,_, rather than cluster densities, from control and detubulated cardiomyocytes. We then addressed the consequences of dystrophin deficiency (*mdx*) and Na_v_1.5 C-terminal truncation (ΔSIV) on Na_v_1.5 expression and organization at the lateral membrane overall, at the groove specifically, and at the T-tubules. ΔSIV resembles the p.V2016M mutation found in a Brugada syndrome patient, replacing the C-terminal valine with a methionine, which results in an apparent reduced expression of Na_v_1.5^5^. ΔSIV knock-in mice display a loss of sodium current and Na_v_1.5 expression specifically at the lateral membrane^5^, and interaction of Na_v_1.5-ΔSIV with the syntrophin/dystrophin complex is impaired^5^. In accordance, dystrophin-deficient *mdx* mice display a strong reduction of sodium current and Na_v_1.5 expression, both overall and specifically at the lateral membrane^7,8^.

By characterizing Na_v_1.5 organization and expression at different pools at a nanoscale and addressing their pool-specific determinants, we come closer to understanding the pathophysiological variety in patients carrying *SCN5A* mutations, as mutations may affect Na_v_1.5 pools and shape the pathophysiological landscape differently. The two genetic mouse models investigated in this study give insight into pool-specific effects of a Na_v_1.5 mutation (ΔSIV) and of disturbing Na_v_1.5 regulation (*mdx*). It appears that (1) Na_v_1.5 is expressed in T-tubules; (2) dystrophin deficiency increases T-tubular Na_v_1.5 expression; and (3) Na_v_1.5 expression is reduced at the lateral membrane of *mdx* and ΔSIV mice overall whereas groove expression is specifically reduced in ΔSIV cells; (4) Na_v_1.5 cluster organization changes at the lateral membrane of *mdx* cells and in the T-tubules of *mdx* and ΔSIV cells compared to wild type.

## RESULTS

### NA_V_1.5 OCCURS AT DISTINCT POOLS IN MURINE VENTRICULAR CARDIOMYOCYTES

The main focus of this work is the characterization of Na_v_1.5 localization and cluster organization in different cardiomyocyte domains with novel quantitative single-molecule localization microscopy (SMLM) and modeling. Technical validation of SMLM is shown in **Supplementary Figure 1**. Firstly, however, we gained a general overview over the Na_v_1.5 pools with confocal microscopy. To this end, we immunostained isolated murine ventricular cardiomyocytes for Na_v_1.5 and the T-tubular marker Bin1. Na_v_1.5 is clearly expressed at the lateral membrane and intercalated disc (**Figure 1c**). Some Na_v_1.5 seems to co-localize with Bin1, the latter showing a striated pattern that indicates the T-tubular membrane. The limited resolution and sensitivity of the confocal microscope however prevents us from establishing a detailed quantitative analysis of Na_v_1.5 in the T-tubules (**Figure 1c**).

To determine the molecular characteristics of Na_v_1.5 expression at the lateral membrane by SMLM, we imaged the lateral membrane of wild-type cardiomyocytes stained for Na_v_1.5 and α-actinin, which coincides with the groove (*N* = 3, *n* = 24; **Figure 2a**). Indeed, Na_v_1.5 occurred both in close proximity to α-actinin and in between α-actinin clusters, indicating localization at the groove and the crest, respectively (**Figure 1a**, **Figure 2a**). To quantify whether the Na_v_1.5 clusters have an affinity for the groove, we used a novel analysis approach^25^ based on Monte-Carlo simulations for each SMLM image (*N* = 3, *n* = 24) in which Na_v_1.5 clusters were redistributed over the image either randomly or with a high affinity for α-actinin (**Figure 2b,c**). The random simulations were performed by selecting the Na_v_1.5 clusters and stochastically redistributing them within the boundaries of the user-defined region of interest (ROI). For high-affinity simulations, an interaction factor of 0.7 was assumed, given that an interaction factor of 0.0 represents random overlap between the two colors, an interaction factor of 1.0 perfect overlap, and an interaction factor equal to or more than 0.5 indicates that one color has significant affinity for the other color^25^. To generate high-affinity simulations with an interaction factor of 0.7, each Na_v_1.5 cluster was first placed randomly within the region of interest, then the program determined whether one or more pixels of the cluster overlapped with an α-actinin cluster. If this overlap was detected, the cluster was kept in place; if not, the program generated a random number between 0 and 1. If that number was greater than 0.7, the cluster was kept in its position; if not, this process was repeated. Overall, these high-affinity simulations generated an image in which Na_v_1.5 clusters overlapped with α-actinin clusters to more than a random degree.

**Figure 2.**
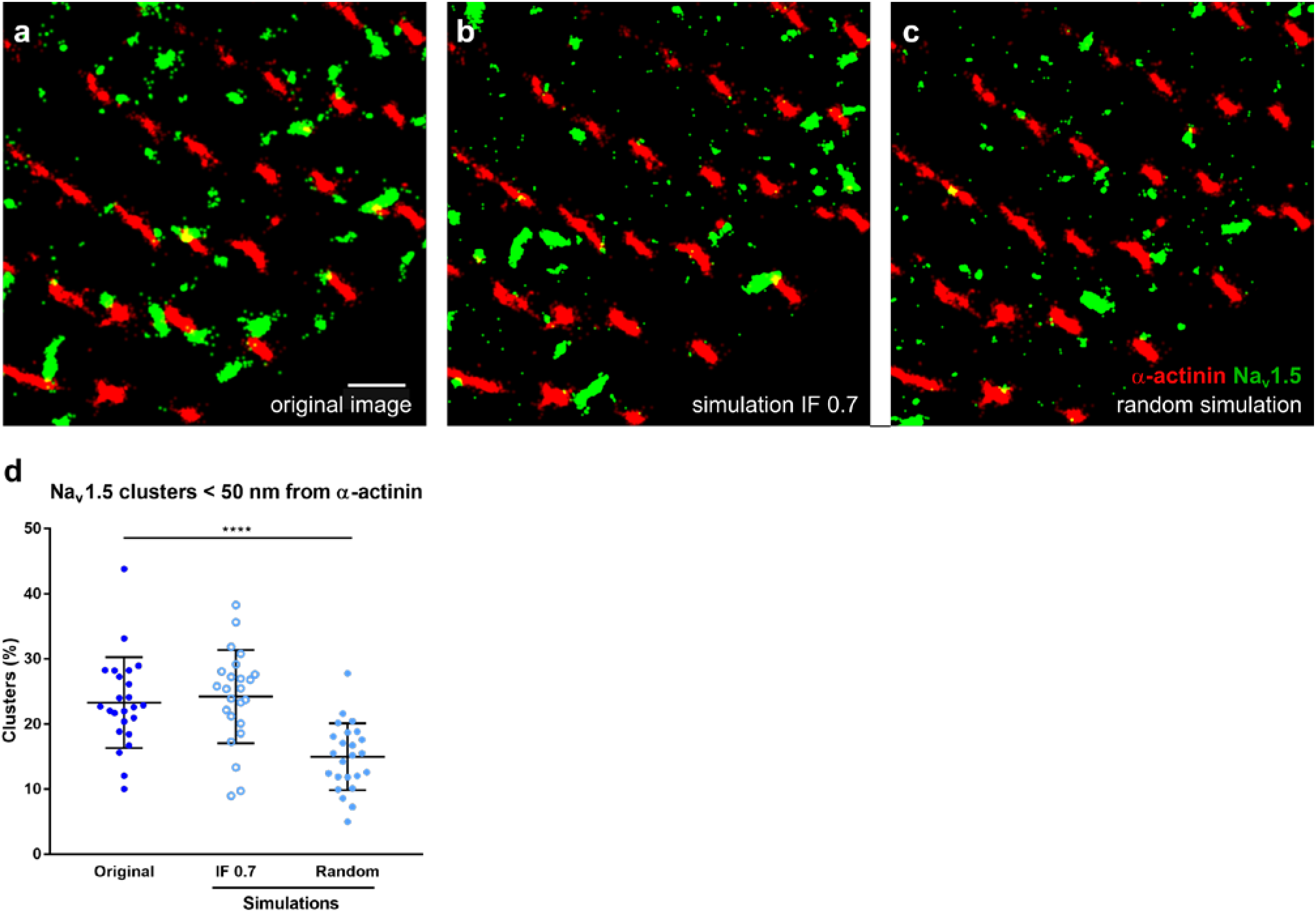
Na_v_1.5 has affinity for the groove at the lateral membrane of a cardiomyocyte. **a**, Super-resolution image of the lateral membrane of murine ventricular myocyte stained for Na_v_1.5 (green) and α-actinin (red). Na_v_1.5 occurs close to α-actinin, indicating the groove; and in between α-actinin lines, indicating the crest. Scale bar 1 μm. **b** and **c** display simulations of the original SMLM image (**a**), in which Na_v_1.5 clusters were randomly distributed (**b**) or had a high affinity for α-actinin (interaction factor [IF] 0.7 [**c**]). Note that differences in affinity are hard to spot by eye, especially in a small area of a cell. **d**, In original SMLM images of wild-type cells (*N* = 3, *n* = 24), the proportion of Na_v_1.5 within 50 nm of α-actinin is ^~^30% higher than in simulations in which Na_v_1.5 clusters of each image are distributed randomly, and similar to images in which a high affinity of Na_v_1.5 for α-actinin was simulated (IF 0.7). ****, *p* < 0.0001, Mann-Whitney test.

Then, for the original and simulated images, the distance from the edge of each α-actinin cluster to the edge of the closest Na_v_1.5 cluster was determined. Na_v_1.5 was considered to be expressed in the groove if it was located within 50 nm from an α-actinin cluster. We found that 25% of Na_v_1.5 clusters were in the groove in experimental images. This value was close to that of the simulations with high affinity for α-actinin (**Figure 2d**), whereas in random simulations, only 15% of Na_v_1.5 clusters were within 50 nm from α-actinin. These results strongly suggest that Na_v_1.5 has an affinity for the groove.

### NA_V_1.5 EXPRESSION AT THE LATERAL MEMBRANE AND THE GROOVE DEPENDS ON DYSTROPHIN AND THE ΔSIV MOTIF, WHEREAS CLUSTER ORGANIZATION DEPENDS ON DYSTROPHIN ONLY

Our SMLM-derived metrics indicate that Na_v_1.5 cluster density at the lateral membrane is reduced by about 30% in *mdx* and ΔSIV cardiomyocytes compared to wild-type cells (**Figure 3a-d**; wild type *N* = 3, *n* = 24; *mdx N* = 3, *n* = 27; ΔSIV *N* = 3, *n* = 19). Cell size and average Na_v_1.5 cluster size did not change between the genotypes, although *mdx* cells showed a high variability in Na_v_1.5 cluster size (**Supplementary Figure 2a**). Interestingly, the solidity and circularity of Na_v_1.5 clusters was markedly increased in *mdx* cells compared to wild type (**Supplementary Figure 2a**). Given that solidity is defined as the ratio between the particle area and the convex hull area of the particle, and circularity as the roundness of a particle (1.0 indicating a perfect circle), the increased solidity and circularity together indicate that Na_v_1.5 clusters are of geometrically less complex shapes. The overall reduction of Na_v_1.5 expression at the lateral membrane in ΔSIV and *mdx* cells additionally confirms the specificity of the anti-Na_v_1.5 antibody since these findings accord with previously reported results^5,7^. In addition, they show that the reduction in whole-cell *I*_Na_ in *mdx* cells is at least partly due to reduced Na_v_1.5 expression at the lateral membrane^8^.

**Figure 3.**
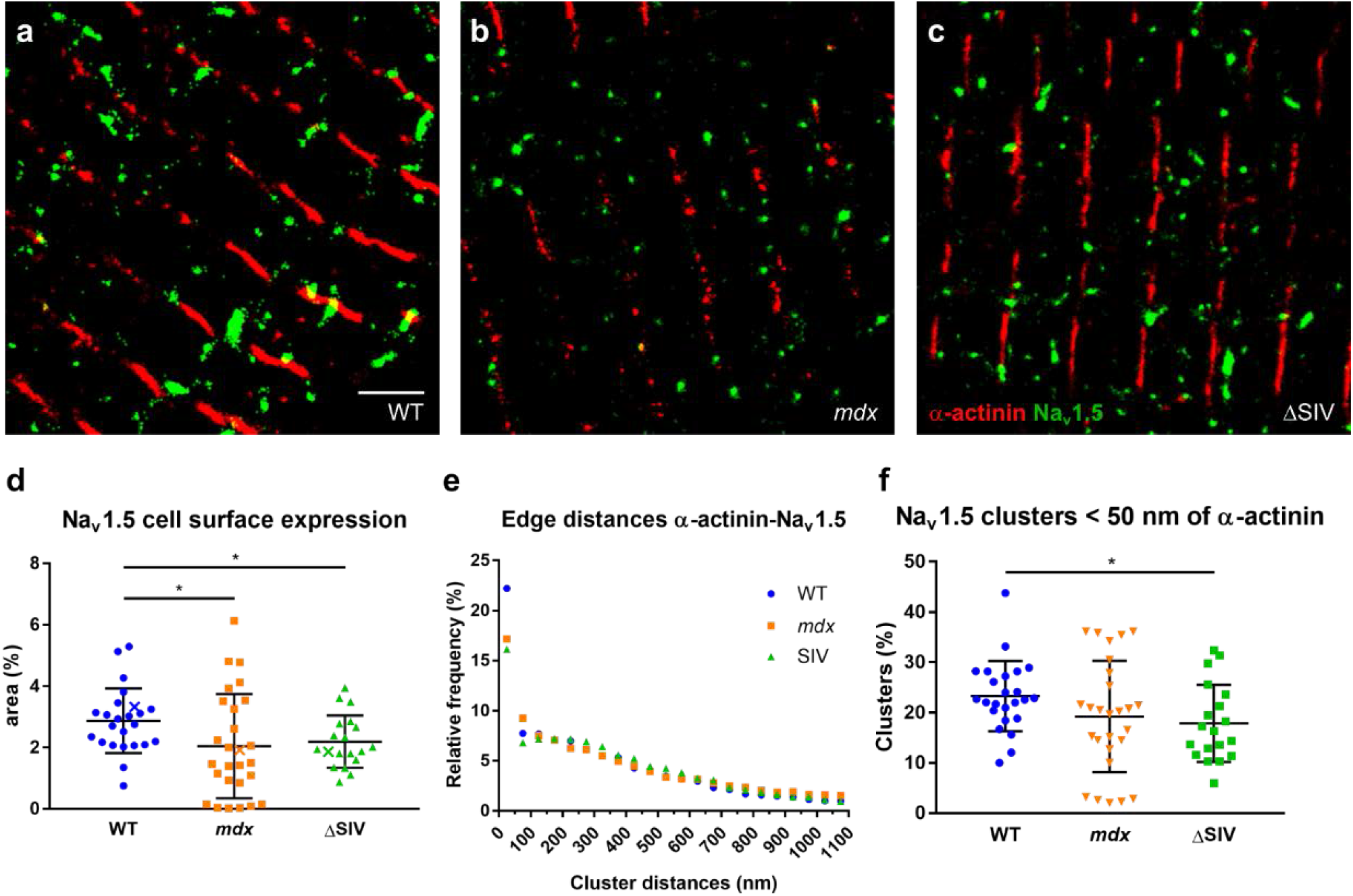
ΔSIV and *mdx* cardiomyocytes show loss of Na_v_1.5 expression at the lateral membrane. **a-c**, Detail of SMLM images of the lateral membrane from wild-type (WT; **a**), *mdx* (**b**), and ΔSIV (**c**) cells. Scale bar 1.5 μm. **d**, Total Na_v_1.5 expression at the lateral membrane is reduced by ^~^30% in *mdx* and ΔSIV compared to wild type. Data points represented by a cross (*X*) correspond to examples from panels **a-c**. **e**, Frequency distribution plot of edge distances from any α-actinin cluster to the closest Na_v_1.5 cluster, bin size 50 nm. In *mdx* and ΔSIV, a smaller proportion of Na_v_1.5 resides within 50 nm to α-actinin compared to wild type. Frequency distributions for cluster distances larger than 50 nm are similar between the genotypes. **f**, Groove expression of Na_v_1.5, defined as Na_v_1.5 clusters within 50 nm of Bin1, is reduced by ^~^30% in ΔSIV cardiomyocytes compared to wild type. Note that *mdx* cells display high variability in Na_v_1.5 cluster expression at the lateral membrane (**d**) and at the groove (**f**). Wild type *N* = 3, *n* = 24; *mdx N* = 3, *n* = 27; ΔSIV *N* = 3, *n* = 19. *, *p* = 0.024 (wt vs. *mdx* [**d**]); *p* = 0.016 (wt vs. ΔSIV [**d**]); *p* = 0.019 (wt vs. ΔSIV [**f**]), Mann-Whitney test.

To assess the effects of Na_v_1.5 truncation and dystrophin deficiency on Na_v_1.5 expression at the crest and the groove, we determined the distances from each α-actinin cluster to the closest Na_v_1.5 cluster in *mdx* and ΔSIV lateral-membrane images, and compared them to wild-type values. The frequency histogram (**Figure 3e**) shows that the proportion of Na_v_1.5 clusters within 50 nm of α-actinin is reduced by ^~^25% in *mdx* and ΔSIV compared to wild-type cells. When the proportion of Na_v_1.5 within 50 nm of α-actinin is plotted for each individual cell (**Figure 3f**), only ΔSIV cardiomyocytes show a significant ^~^20% reduction compared to wild type. Although mean Na_v_1.5 expression in *mdx* cells is reduced by about 17% compared to wild type, the data show high variability (**Figure 3f**), which correlates with the high variability in Na_v_1.5 expression at the lateral membrane of *mdx* cells (**Figure 3d**). The affinity of Na_v_1.5 for the groove is however similar in wild-type, *mdx*, and ΔSIV cells when comparing the proportion of Na_v_1.5 clusters within 50 nm of α-actinin from experimental images, and random and high-affinity simulations for each genotype (**Supplementary Figure 3a**).

Taken together, these results establish that (1) Na_v_1.5 expression at the groove of the lateral membrane partially depends on the ΔSIV motif and dystrophin expression; (2) in dystrophin-deficient cells, Na_v_1.5 clusters at the lateral membrane are organized into simpler shapes compared to wild-type cells; and (3) dystrophin deficiency induces high variability in Na_v_1.5 expression and Na_v_1.5 cluster size.

### NA_V_1.5 IS EXPRESSED IN THE T-TUBULAR MEMBRANE

Na_v_1.5 expression in the T-tubules was assessed by co-staining cardiomyocytes for Na_v_1.5 and the T-tubular marker Bin1, and imaging an intracellular plane of each cell in highly inclined and laminated optical (HILO) mode. Bin1 was chosen as a T-tubular marker since it binds and shapes the T-tubular membrane and assists in the trafficking and clustering of the voltage-gated calcium channel Ca_v_1.2^24,26^. The Bin1 antibody was first validated by confirming that Bin1 stainings are in close proximity to α-actinin, given that T-tubules run in close proximity to the sarcomeric Z-disc (*N* = 3, *n* = 59) (**Supplemental figure 4**)^13^. Then, we assessed whether Na_v_1.5 and Bin1 associated randomly or not by performing simulations for each SMLM image (*N* = 3, *n* = 39), where Na_v_1.5 clusters were redistributed either randomly or with high affinity for Bin1 (**Figure 4a-c**). Na_v_1.5 clusters at the lateral membrane and intercalated disc were excluded. Then, the distance from each Bin1 cluster to the closest Na_v_1.5 cluster was determined. The frequency histogram shows that about 16-17% of Na_v_1.5 clusters is within 50 nm of Bin1 in both original images and high-affinity simulations, whereas in random simulations this value was only ^~^8% (**Figure 4d**). This was confirmed when plotting the proportion of Na_v_1.5 clusters within 50 nm of Bin1 for each individual image (**Figure 4e**). Together, these findings indicate that a considerable subset of intracellular Na_v_1.5 is expressed in the T-tubular membrane.

**Figure 4.**
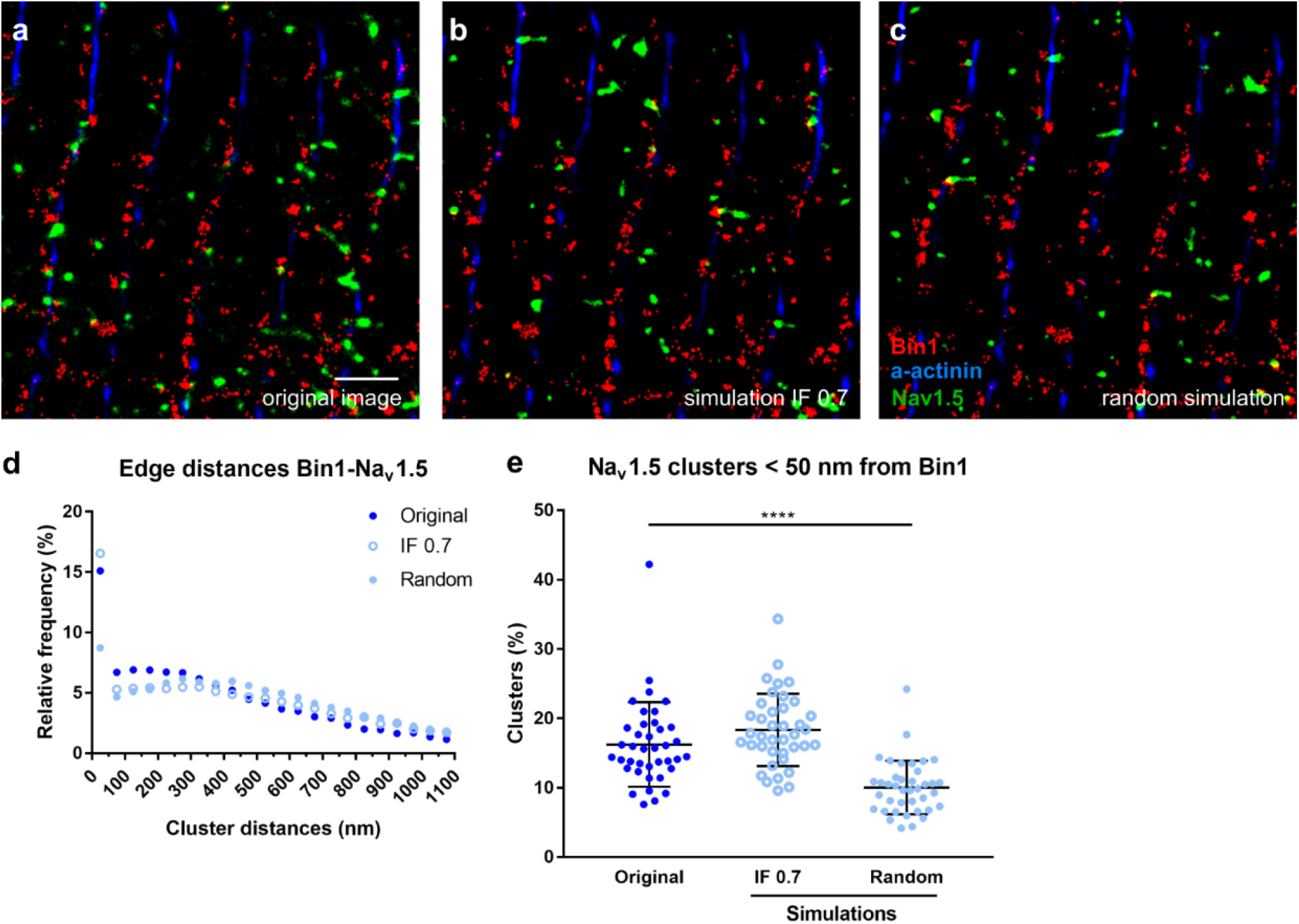
Na_v_1.5 is expressed in the T-tubules. **a**, Detail of an intracellular recording from a wild-type cardiomyocyte stained for α-actinin (blue), Bin1 (red), and Na_v_1.5 (green). All Na_v_1.5 clusters are redistributed over the image either randomly (**b**) or with a high affinity for Bin1 (interaction factor [IF] 0.7 [**c**]). Scale bar 1.5 μm. **d**, For each Bin1 cluster, the closest Na_v_1.5 cluster is within 50 nm in ^~^15% of cases, which is significantly more than the ^~^8% from random simulations and close to ^~^17% of the high-affinity simulations (IF 0.7) (*N* = 3, *n* = 39). The frequency distributions of Na_v_1.5 for distances to α-actinin of more than 50 nm does not show any apparent difference between the genotypes. **e**, In experimental SMLM images, a higher proportion (^~^40%) of Na_v_1.5 are within 50 nm from the nearest Bin1 cluster than in random simulations, but not as much as in simulations with a high-affinity simulations (IF 0.7). Values for each individual cell/simulation are plotted. ****, *p* < 0.0001, unpaired t-test.

### T-TUBULAR SODIUM CURRENT CANNOT BE ASSESSED BY WHOLE-CELL RECORDINGS

Having shown that Na_v_1.5 is expressed at the T-tubules (**Figure 4**), we investigated whether we could record a T-tubular sodium current. To this end, we compared whole-cell *I*_Na_ recordings from normal (*N* = 5, *n* = 7) and detubulated (N = 5, *n* = 7) ventricular cardiomyocytes. Firstly, we concluded that detubulation was successful as the capacitance of detubulated cells was 30% lower than of control cells (**Figure 5b**). Whole-cell *I*_Na_ did not decrease after detubulation (**Figure 5a**), which indicates that the vast majority of sodium current is conducted by sodium channels at the intercalated disc and lateral membrane. Comparing the current density shows a 25% yet non-significant increase in *I*_Na_ density after detubulation (**Figure 5c**), further confirming that the majority of sodium channels is outside the T-tubular domain. As discussed in the introduction, current densities of detubulated and control cardiomyocytes cannot be compared directly as capacitance measurements in control cardiomyocytes result in a considerable underestimation of cell membrane area. Thus, our data show that the T-tubular *I*_Na_ is below the detection limit of the whole-cell patch-clamp technique.

**Figure 5.**
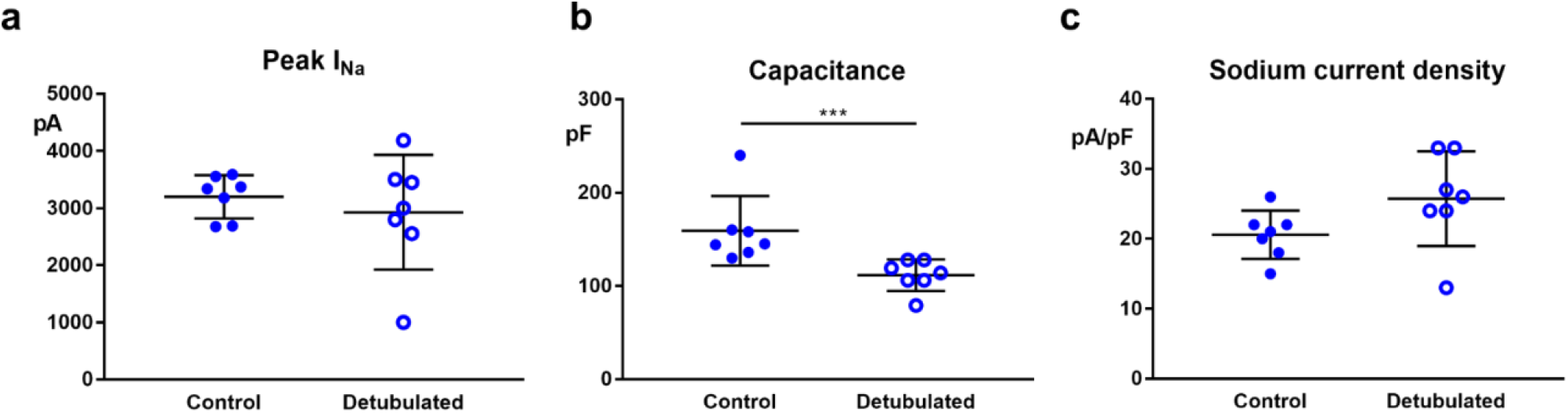
Whole-cell sodium current recordings in detubulated and normal cardiomyocytes. Compared to control cells, maximum sodium current (**a**) didn’t change, capacitance (**b**) was reduced by about 30%, and sodium current density (**c**) increased by about 25% (non-significant) in detubulated cells. *N* = 5, *n* = 7 for both conditions. ***, *p* = 0.006, Mann-Whitney test.

### NA_V_1.5 EXPRESSION IS INCREASED IN T-TUBULES FROM DYSTROPHIN-DEFICIENT CARDIOMYOCYTES, AND CLUSTER ORGANIZATION DEPENDS ON THE SIV MOTIF AND DYSTROPHIN

We next investigated whether dystrophin deficiency (*mdx*) and deletion of the SIV motif of Na_v_1.5 affected Na_v_1.5 expression in the T-tubules. Firstly, we determined that overall Na_v_1.5 cluster density in intracellular planes was increased in *mdx* compared to wild-type cells, whereas no difference was observed between ΔSIV and wild-type cells (**Figure 6a**; wild type *N* = 3, *n* = 47; *mdx N* = 3, *n* = 26; ΔSIV *N* = 3, *n* = 13). We also noted an increase in cluster solidity in *mdx* and an increase in cluster circularity in *mdx* and ΔSIV compared to wild-type cells, while cell size and average Na_v_1.5 cluster size did not differ between the three genotypes. Similar to what we observed at the lateral membrane, this indicates that intracellular Na_v_1.5 clusters have geometrically simpler shapes in *mdx* and ΔSIV cells.

**Figure 6.**
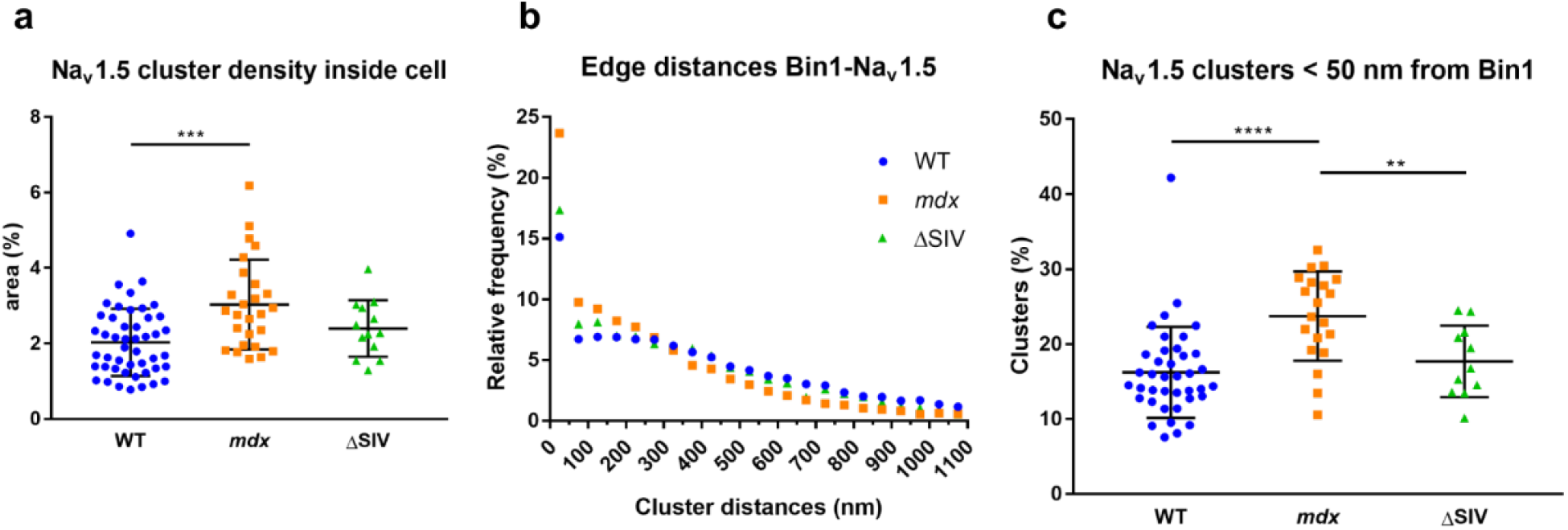
Na_v_1.5 expression is increased in T-tubules of *mdx* mice. **a**, Na_v_1.5 cluster density in intracellular membrane compartments and T-tubules is ^~^30% higher in *mdx* cardiomyocytes than in wild-type cardiomyocytes, but no change was observed in ΔSIV compared to wild-type cells. **b**, For each Bin1 cluster, the distance to the closest Na_v_1.5 cluster is measured. Cluster distances are plotted in a frequency histogram with bins of 50 nm. The given histogram indicates that Na_v_1.5 preferentially locates within 50 nm to Bin1, in ΔSIV and *mdx* more so than in wild-type cells. **c**, In *mdx* cells, a higher proportion of Na_v_1.5 clusters is within 50 nm from Bin1 than in wild-type cells. Together with the increase in overall intracellular Na_v_1.5 expression in *mdx*, this indicates that T-tubules in *mdx* cells show a higher Na_v_1.5 expression. ΔSIV cells show a similar T-tubular Na_v_1.5 expression as WT cells. **a**, Wild type *N* = 3, n = 47; *mdx N* = 3, *n* = 26; ΔSIV *N* = 3, *n* = 13. **b,c** Wild type *N* = 3, n = 39; *mdx N* = 3, *n* = 20; ΔSIV *N* = 3, *n* = 11. **, *p* = 0.0095; ***, *p* = 0.0002; ****, *p* < 0.0001, Mann Whitney test.

The frequency histogram plotting distances between Bin1 and Na_v_1.5 shows that the proportion of Na_v_1.5 within 50 nm to Bin1 was ^~^30% higher in *mdx* and ^~^4% higher in ΔSIV compared to wild type (**Figure 6b**; wild type *N* = 3, n = 39; *mdx N* = 3, *n* = 20; ΔSIV *N* = 3, *n* = 11). Plotting these values from individual cells confirmed this increase of T-tubular Na_v_1.5 in *mdx* cells, but no difference between ΔSIV and wild type cells was observed (**Figure 6c**). When comparing T-tubular Na_v_1.5 expression from original images and random and high-affinity simulations, Na_v_1.5 displays an affinity for Bin1 in all three genotypes; but in *mdx* cells, Na_v_1.5 has an even higher affinity for Bin1 than in the high-affinity simulations (**Supplementary figure 3b**).

Taken together, these data indicate that dystrophin deficiency induces a higher Na_v_1.5 expression associated with intracellular membrane compartments and T-tubules compared to wild type, whereas this effect was not observed in ΔSIV cells. Intracellular Na_v_1.5 cluster shapes were also markedly simpler in ΔSIV and *mdx* compared to wild-type cardiomyocytes.

## DISCUSSION

This work aimed to surpass the previously published limited-resolution characterization of Na_v_1.5 expression and cluster organization at the lateral membrane and T-tubules of cardiomyocytes. To this end, we applied novel quantitative single-molecule localization microscopy (SMLM) and modeling techniques to investigate Na_v_1.5 organization at the crest, groove, and T-tubules in cardiomyocytes from wild-type mice, dystrophin-deficient (*mdx*) mice, and mice expressing C-terminally truncated Na_v_1.5 (ΔSIV). We showed that (1) Na_v_1.5 expression in the groove of the lateral membrane partly depends on dystrophin and the SIV motif of Na_v_1.5; (2) dystrophin is involved in Na_v_1.5 cluster organization at the lateral membrane; (3) Na_v_1.5 is expressed in the T-tubules, although we could not assess T-tubular sodium current with our electrophysiological approach; (4) T-tubular Na_v_1.5 expression is increased in dystrophin-deficient but not in ΔSIV cardiomyocytes; and (5) intracellular Na_v_1.5 cluster organization depends on dystrophin and the SIV motif. These findings are schematically summarized in **Figure 7**.

**Figure 7.**
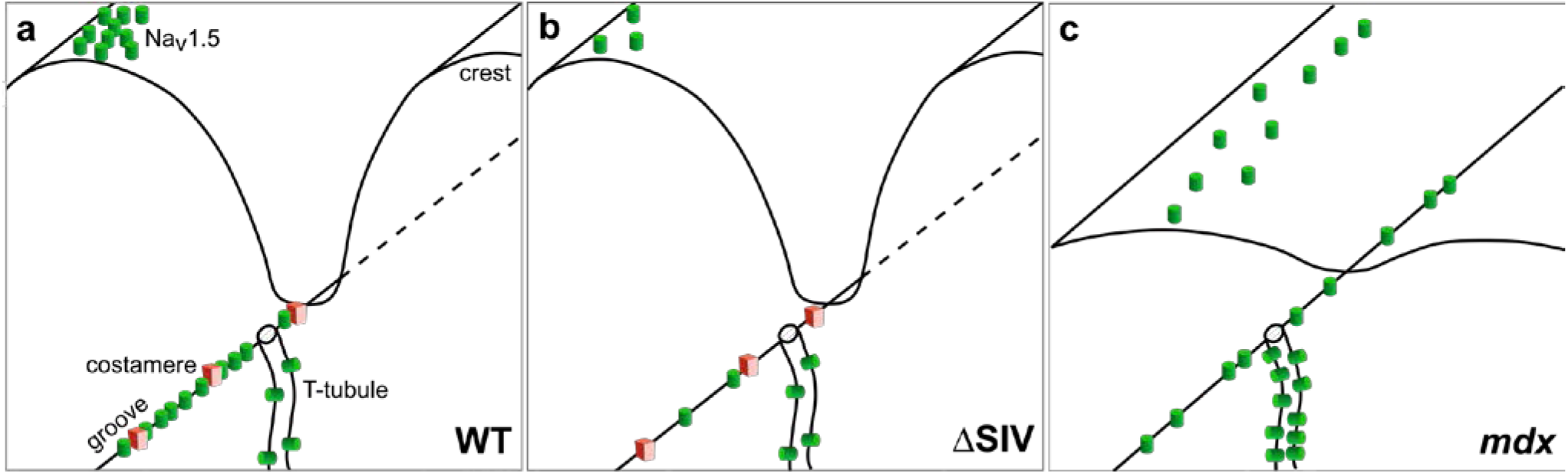
Model depicting changes at the lateral membrane and T-tubules of ΔSIV and *mdx* cardiomyocytes. Compared to the wild-type condition (**a**), ΔSIV cells (**b**) show less Na_v_1.5 at the lateral membrane overall and specifically in the groove, whereas the lateral membrane structure remain intact. In *mdx* cells (**c**), the lateral membrane is flatter than in wild-type and ΔSIV cells, and Na_v_1.5 expression at the lateral membrane is reduced overall; yet the expression at groove of *mdx* cells is not as affected as that of ΔSIV cells. Costameres are severely compromised in *mdx* cells. In the T-tubules, Na_v_1.5 expression is similar in wild-type (**a**) and ΔSIV cells (**b**), and increased in *mdx* cardiomyocytes (**c**).

We convincingly showed that Na_v_1.5 is expressed in the T-tubules by co-staining Na_v_1.5 with the T-tubular marker Bin1 in SMLM recordings, and assessing the robustness of this association by novel *in silico* methods^25,27^. Comparing original images to simulations of these images in which Na_v_1.5 clusters were redistributed either randomly or with high affinity for Bin1, we observed that the T-tubular population of Na_v_1.5 in original images is twice as high as in random populations and comparable to that from high-affinity simulations (**Figure 4e**). The function of Na_v_1.5 at the T-tubules remains elusive, although Na_v_1.5 in the T-tubules will theoretically increase conduction velocity^28^. Since some Bin1 and Na_v_1.5 clusters overlapped (**Figure 4a**), we may hypothesize that Bin1 could be involved in the regulation of Na_v_1.5 in the T-tubules as Bin1 also regulates the trafficking of voltage-gated calcium channel Ca_v_1.2^29^.

We showed that the presence of a T-tubular *I*_Na_ could not be deducted from whole-cell electrophysiological methods as whole-cell currents between detubulated and control cardiomyocytes were similar (**Figure 5a**). The T-tubular *I*_Na_ may not surpass the noise of whole-cell recordings, or the T-tubular membrane may not be properly voltage clamped in our whole-cell approach, thus a proportion of T-tubular sodium channels may not be activated (space clamp issue). In the past, the T-tubular *I*_Na_ was suggested to be ^~^30% of total *I*_Na_ by comparing current densities from detubulated and control cells^16^. Measuring the capacitance in control cells underestimates cell membrane area, however, which disqualifies the direct comparison of current densities non-detubulated to detubulated cells^17^. Other electrophysiological methods have not been able to record T-tubular currents either: cell-attached^30^ and SICM data^11,31^ only show currents from a few voltage-gated calcium channels whereas a much stronger calcium current would be expected from the T-tubules. To assess the T-tubular *I*_Na_, more specific methods need to be developed.

When dystrophin is absent, less Na_v_1.5 is expressed at the lateral membrane (**Figure 3a**), but more Na_v_1.5 is expressed in the T-tubules compared to wild-type cells (**Figure 6c**). The reduction Na_v_1.5 at the lateral membrane of *mdx* cells may be explained by the loss of costamere integrity (**Figure 7**)^32^. We may also hypothesize that dystrophin deficiency causes a partial rerouting of Na_v_1.5 trafficking to the T-tubules instead of to the lateral membrane by an unknown mechanism. The short dystrophin product Dp71 may be involved since it is present only at the T-tubules^33^, and binds ion channels and the cytoskeleton^34^. In light of the overall reduction of Nav1.5 protein expression and *I*_Na_ in *mdx* hearts^8^, the relative increase of T-tubular Na_v_1.5 expression may make matters worse: due to the restricted space in the T-tubular lumen, the T-tubular sodium current may partially self-attenuate^6,35^. A *de facto* translocation of Na_v_1.5 from the lateral membrane to the T-tubules of *mdx* cells may quench whole-cell sodium current and contribute to the reduction in ventricular conduction velocity observed in *mdx* mice^6,8^. Whether this effect indeed takes place *in vivo* requires further research.

Interestingly, Na_v_1.5 cluster shapes in the groove and the intracellular compartment are simpler in *mdx* than in wild-type cells, illustrated by the increased solidity and circularity (**Supplementary Figure 2**). This indicates that dystrophin is involved in scaffolding and shaping of the Na_v_1.5 clusters, which may be related to the large size (427 kD) of dystrophin, and to the link to the actin cytoskeleton that dystrophin provides^36^. On the other hand, Na_v_1.5 clusters in ΔSIV cells show an increase in solidity only in intracellular planes (**Supplementary Figure 2b**). This suggests that the interaction of Na_v_1.5 with the syntrophin-dystrophin plays a role in cluster organization in the intracellular compartment/T-tubules, while at the lateral membrane, the secondary N-terminal syntrophin-binding site of Na_v_1.5 may support interaction with dystrophin and wild-type-like Na_v_1.5 cluster organization^37^.

The residual Na_v_1.5 expression at the lateral membrane of *mdx* mice may be explained by the dystrophin homologue utrophin^38^. Utrophin is expressed during the fetal phase and re-expressed in adult tissue when dystrophin is absent in mice, but not in humans^38,39^. Since we found that Na_v_1.5 cluster density and size at the lateral membrane and at the groove is highly variable in *mdx* cells (**Figure 3a,c, Supplementary Figure 2a**), ventricular cardiomyocytes may display a high variability in compensatory utrophin and Dp71 expression. Utrophin however does not rescue a wild-type-like phenotype: the costameres and sarcolemma of utrophin-expressing *mdx* mice are still weak^40^, and the crest-groove profile at the lateral membrane of *mdx* cardiomyocytes is flattened compared to wild type (**Figure 7c**)^41^. The effects of dystrophin deficiency when the aforementioned compensatory mechanisms are not available needs to be assessed in heart samples from Duchenne patients and from dystrophin/utrophin/Dp71 triple knock-out mice.

Like *mdx* mice, ΔSIV mice express less Na_v_1.5 at the lateral membrane, and the groove pool is specifically affected (**Figure 3d,f**). The ΔSIV truncation reduces the interaction of Na_v_1.5 with the syntrophin-dystrophin complex^5^. Since ΔSIV mice still express dystrophin, costameres presumably remain intact. Of note, the syntrophin-dystrophin complex may also interact with the N-terminus of Na_v_1.5^37^, possibly explaining the reduction, rather than the abolition, of Na_v_1.5 localization to the groove and the lateral membrane. Contrary to *mdx* mice, we did not observe an increase of Na_v_1.5 at the T-tubules in ΔSIV mice, which may be explained by their intact costameres.

A groove-specific reduction of Na_v_1.5, as shown for ΔSIV and a subset of *mdx* cardiomyocytes, and an increase in T-tubular Na_v_1.5, as shown for *mdx* cells, may affect conduction and excitability in different ways. However, late openings between the crest and the groove as well as biophysical properties did not differ in murine cardiomyocytes^11^. Future studies should address specific differences in function and regulation of Na_v_1.5 in the crest, groove, and T-tubules.

Intriguingly, while our data revealed that Na_v_1.5 is reduced at the groove of *mdx* and ΔSIV mice, Rivaud *et al*. have recently shown in a murine heart failure model (transient aortic constriction) that sodium current is specifically reduced at the crest, not at the groove, compared to sham-operated animals^11^. Rivaud *et al*. showed that Na_v_1.5 cluster size, not cluster density, was markedly reduced at the lateral membrane in the heart failure model, whereas our data show that cluster density, not cluster size, was reduced in *mdx* and ΔSIV cells (**Supplementary Figure 2a**). Expression and organization of Na_v_1.5 at the crest and the groove was however not investigated in this heart failure model. Together, these data suggest that cardiomyocytes respond differently to dystrophin deficiency, Na_v_1.5 truncation (ΔSIV), and transient aortic constriction^11^. One obvious difference is the hypertrophy observed in the heart failure model but not in *mdx* or ΔSIV mice. These different pathways remain to be elucidated in detail.

The phenotype of Brugada syndrome patients carrying the mutation p.V2016M, which changes the C-terminal motif SIV into SIM, may be partly explained by a reduction in Na_v_1.5 cluster density in the groove of the lateral membrane and a reduction of dystrophin-Na_v_1.5 interaction^8^. Indeed, in ΔSIV cardiomyocytes, transversal conduction velocity, total sodium current, and sodium current at the lateral membrane are decreased^5^, which may contribute to the Brugada syndrome phenotype. However, the specific effect of groove-specific Na_v_1.5 reduction remains to be determined.

A limitation of our study is that *mdx* mice display various compensatory mechanisms discussed above that do not allow us to assess the effects of a “pure” dystrophin deficiency. *Mdx* mice moreover do not have the optimal controls, since they were on a Bl6/Ros background while being backcrossed to Bl/6J in the fourth generation, whereas wild-type and ΔSIV mice were littermates on a pure Bl6/J background. The discrepancy in backgrounds may have introduced variability in our data, although the data are consistent with previously published results^7,8^. Lastly, our methods did not allow us to quantify crest expression of Na_v_1.5.

Taken together, our findings provide important and previously unattainable mechanistic insights on the properties of Na_v_1.5 organization on the crest, groove, and lateral membrane of cardiomyocytes, and are an important step towards identifying cardiac domain-specific molecular determinants of Na_v_1.5 and location-specific effects of *SCN5A* mutations. However, pool-specific function and regulation of Na_v_1.5 at the crest, groove, and T-tubules remains to be studied in more detail.

## METHODS

### ETHICAL STATEMENT

All animal experiments conformed to the *Guide to the Care and Use of Laboratory Animals* (US National Institutes of Health, publication No. 85-23, revised 1996); were approved by the Cantonal Veterinary Administration, Bern, Switzerland; conformed to the New York University guidelines for animal use and care (IACUC Protocol 160726-03 to MD, approved 07/11/2018); and have complied with the Swiss Federal Animal Protection Law.

### MOUSE MODELS

#### Dystrophin knock-out (*mdx*^5CV^) mice

The *mdx^5CV^* mouse strain displays total deletion of dystrophin protein. It was generated as described previously^42^, and purchased from the Jackson laboratory (Bar Harbor, Maine; stock #002379). *Mdx* mice were on a mixed background of Bl6/Ros and Bl6/J (fourth generation of backcrossing). Male littermates aged 16 weeks were used.

#### ΔSIV knock-in mice

In ΔSIV knock-in mice, the C-terminal SIV motif of Na_v_1.5 is deleted (*Scn5a*-p.S2017STOP), as described previously^5^. ΔSIV mice and wild-type littermates are on a pure Bl6/J background. All mice (male and female) were 16 weeks old.

### ISOLATION OF MURINE VENTRICULAR MYOCYTES

All experimental steps are performed at room temperature unless specified otherwise.

Cardiomyocytes were isolated based on a previously established enzymatic method^8^. Briefly, mice were euthanized by cervical dislocation. Hearts was excised, cannulated, and mounted on a Langendorff system for retrograde perfusion at 37°C. Hearts were rinsed first with nominally Ca^2+^-free solution containing (in mM) 135 NaCl, 4 KCl, 1.2 MgCl_2_, 1.2 NaH_2_PO_4_, 10 HEPES, and 11 glucose (pH 7.4, NaOH adjusted), then with digestion solution for 15 minutes, consisting of the aforementioned solution supplemented with 50 μM Ca^2+^ and collagenase type II (1 mg/mL; Worthington, Allschwil, Switzerland). Subsequently, atria were removed. Ventricles were transferred to nominally Ca^2+^-free solution and minced into small pieces. To increase the yield of isolated cells, digested ventricular tissue was triturated gently and filtered through a 100 μm nylon mesh. Before use, cells were subjected to a calcium increase procedure.

### CONFOCAL MICROSCOPY

Isolated cardiomyocytes were fixed with acetone at −20°C for 10 minutes in cell culture chambers. Cells were washed twice with phosphate-buffered saline (PBS), permeabilized with 0.1% saponin in PBS for 2 x 7 minutes, and blocked for 15 minutes with PBS containing 10% normal goat serum, 1% bovine serum albumin (BSA), 0.1% saponin, and 30 μg/mL Fab fragment donkey anti-mouse IgG (H+L) (Jackson Immuno Research, Baltimore, Maryland). All following antibody dilutions and washing steps were performed with incubation buffer (3% normal goat serum, 0.05% saponin, and 1% BSA in PBS) unless specified otherwise. Primary antibodies were diluted and applied onto cells for overnight incubation at 4°C. Afterwards, cells were washed 4 times for 5 minutes, and incubated with secondary antibodies for 2 hours. Cells were washed 3 times for 5 minutes, incubated with DAPI (1 μL in 200 μL PBS) for 20 minutes, washed twice with PBS, and finally embedded in FluorSave reagent (Merck, Burlington, Massachusetts). Cells were imaged on a confocal microscope (LSM710, Zeiss, Oberkochen, Germany).

### SINGLE-MOLECULE LOCALIZATION MICROSCOPY

#### Sample preparation

Freshly isolated cardiomyocytes were plated on laminin-coated glass coverslips and allowed to adhere at 37°C for 30 minutes. Cells were fixed with 4% paraformaldehyde for 10 minutes and washed 3 times with PBS. To quench autofluorescence, cells were placed overnight under 470 nm LED light. Then, cells were permeabilized with 0.1% Triton X-100 in PBS for 10 minutes, and blocked with blocking solution (2% glycine, 2% BSA, and 0.2% gelatin in PBS) for 30 minutes. All following antibody dilutions and washes were performed with blocking solution unless specified otherwise. Unconjugated primary antibodies were applied to the cells for 1 hour, followed by three 5-minute washes. Secondary antibodies were applied for 15 minutes, followed by three 5-minute washes and an optional 15-minute incubation with conjugated primary antibodies. Lastly, cells were washed with PBS for 5 minutes. Coverslips were mounted onto microscope slides in which two small holes were drilled and using double-sided tape as spacers, creating a fluid chamber. Chambers were sealed with epoxy resin.

#### Optical setup and image acquisition

SMLM imaging was performed in accordance to a previously described method^23^. To achieve stochastic fluorophore blinking, imaging buffer was added to the cells, containing 200 mM 2-mercaptoethylamine, 10% glucose, 0.04 mg/mL glucose oxidase and 0.08 mg/mL catalase in T50 buffer (10 mM Tris pH 8 and 50 mM NaCl in MilliQ). Samples were imaged on a customized Leica DMI 300 inverse microscope equipped with an HCX PL APO 63X NA = 1.47 OIL CORR TIRF objective (Zeiss), a 2X tube lens (Diagnostic Instruments, Sterling Heights, Michigan) and a chromatic aberration correction lens (AC254-300-A; Thorlabs, Newton, New Jersey). Samples were sequentially excited by a 639 nm laser (, MRL-FN-639-800; UltraLasers, Newmarket, Canada), 561 nm laser (MGL-FN-561-200; UltraLasers), and 488 nm laser (OBIS; Coherent, Santa Clara, California). A 405 nm laser (MDL-III-405-150; CNI, Changchun, China) was used to reactivate AlexaFluor 647 fluorophores. Lasers were aligned by a penta-edged dichroic beam splitter (FF408/504/581/667/762-Di01-22×29). The 488, 561, and 639 laser lines were adjusted to ~0.8, 1.0, and 1.5 kW/cm^2^, respectively. The emitted fluorescence was filtered by the applicable single-band fluorescence filter (FF01-531/40, FF01-607/36, and FF01-676/37 for AlexaFluor 488, 568, and 647, respectively; Semrock, Rochester, New York) in a filter wheel, and recorded at 33 Hz by a sCMOS camera (Prime95B; Photometrics, Tucson, Arizona) with a minimum of 2000 frames per wavelength. Recordings were controlled by MicroManager software (version 1.4.22; www.micro-manager.com). The readout noise of each camera pixel was pre-calibrated and characterized by a Gaussian distribution^43^. Signals from the lateral membrane were recorded in the first focal plane that contained α-actinin signal having moved the focal plane up from the cover glass. Intracellular/T-tubular signals were recorded in a higher focal plane with the TIRF objective in HILO (highly inclined and laminated optical) mode.

#### Alignment of different colors for SMLM images

Different colors were aligned following previously described methods^23^. Briefly, broad-spectrum fluorescent beads (diameter ~100 nm, TetraSpec; Thermo Fisher, Waltham, Massachusetts) were imaged sequentially in the blue (488 nm), green (568 nm), and red (639 nm) channels. The images from the blue and green channels were aligned to the red channel using a second polynomial warping algorithm in Matlab (version R2017b; Mathworks, Natick, Massachusetts).

#### Single-molecule localization

Single-molecule localization also followed a previously described method^23^. In short, from each image stack, each frame was box-filtered. Box size was four times the full width at half maximum of a 2D Gaussian point spread function (PSF). Each pixel was weighted by the inverse of its variation during such box-filtering. The low-pass filtered image was extracted from the raw image, and local maxima were identified. Local maxima from all frames of one image stack were subjected to 2D Gaussian multi-PSF fitting^44^. The 2D Gaussian single-PSF fitting was performed using the maximum likelihood estimation (MLE) algorithm on a GPU (GTX 1060, CUDA 8.0; Nvidia, Santa Clara, California). The likelihood function at each pixel was based on convolving the Poisson distribution of the shot noise, which is governed by the photons emitted from nearby fluorophores, and the Gaussian distribution of the readout noise characterized by the pre-calibrated expectation, variation, and analog-to-digital conversion factor. Fitting accuracy was estimated by Cramér-Rao lower bound.

#### Image analysis

Images were processed based on previously described methods^27^. Briefly, images were subjected to a smoothing filter, adjusted for brightness and contrast, and filtered to a threshold to obtain a binary image with ImageJ software (version 1.51j8; https://imagej.net). Regions of interest were drawn, defined as regions with regular α-actinin and/or Bin1 staining, depending on the applied antibodies. We excluded the intercalated disc regions from both surface and intracellular recordings, and the lateral membrane regions from intracellular recordings. Clusters were analyzed in ImageJ. Image simulations were performed using the Interaction Factor plugin in ImageJ^25^, and distance between clusters in experimental and simulated images were obtained using the ImageJ function ‘Analyze particles’ and a script written in Python (version 2.7; www.python.org) that utilized the image processing packages ‘scikit-image’ and ‘mahotas’^27^. Frequency histograms of cluster distances were made in GraphPad Prism (version 7; GraphPad Software, San Diego, CA, USA). The investigator that analyzed the images was ignorant of the corresponding genotypes.

### ANTIBODIES

Primary antibodies were custom-made rabbit anti-Na_v_1.5 (epitope: amino acids 493-511 of rat Na_v_1.5; Pineda Antibody Service, Berlin, Germany; 1:100), mouse anti-Bin1 (amphiphysin-II; epitope: amino acids 179-207 of human Bin1; Santa Cruz Biotechnology, Dallas, Texas; 1:50), mouse anti-α-actinin (raised against rabbit skeletal α-actinin; Sigma-Aldrich, St. Louis, Missouri; 1:400), and the same anti-α-actinin antibody directly conjugated to AlexaFluor 488 or 647 (Thermo Fisher; 1:2000 or 1:5000, respectively). Secondary antibodies were rabbit AlexaFluor 568 (Life Technologies, Carlsbad, California; SMLM: 1:10,000; confocal: 1:200), mouse AlexaFluor 647 (Life Technologies; SMLM: 1:5000), and mouse AlexaFluor 488 (Life Technologies; confocal: 1:200).

#### Antibody specificity

The specificity of the Pineda rabbit anti-Na_v_1.5 antibody was determined as follows. Firstly, the confocal image in **Figure 1** shows that the staining pattern of the Pineda antibody is highly similar to that of the widely used Alomone ASC-005 antibody^5^. Secondly, we have previously published proximity ligation assay data using the Pineda antibody that show a loss of interaction between Na_v_1.5 and α-syntrophin at the lateral membrane in ΔSIV mice compared to wild type. This corresponds with the lateral-membrane-specific reduction of *I*_Na_ and Na_v_1.5 staining using the Alomone antibody in ΔSIV mice compared to wild type^5^. Our group has shown a similar reduction of Na_v_1.5 at the lateral membrane of *mdx* mice using the aforementioned Alomone antibody^7^. Our SMLM data generated using our Pineda antibody reproduced this reduction of Na_v_1.5 expression at the lateral membrane in ΔSIV and *mdx* mice (**Figure 3D**). Moreover, the Na_v_1.5 localization patterns in our SMLM data generated using the Pineda antibody shows great similarities with those generated using a conventional Sigma (S0189) anti-Na_v_1.5 antibody^45^.

Lastly, we showed that the Pineda Na_v_1.5 antibody does not recognize other relevant Na_v_ isoforms (**Supplemental Figure 5**). We stained human embryonic kidney (TsA-201) cells transiently expressing Na_v_1.5 or Na_v_1.4 with the Pineda antibody, because, from all Na_v_-encoding genes, only relevant amounts of Na_v_1.5- and Na_v_1.4-encoding mRNA are expressed in cardiomyocytes from the mice used in our study^46^. Briefly, TsA-201 cells (ATCC, VA, USA) were cultured in Dulbecco ‘ s modified Eagle’s culture medium (Gibco, Thermo Fisher Scientific, MA, USA) supplemented with 2 mM glutamine and 10% heat-inactivated fetal bovine serum, at 37°C with 5% CO_2_. Cells were transfected with the following vectors: pEGFP-c1 and pcDNA3.1-FLAG-*hSCN5A*-WT, pRC/CMV-hSkM1=Nav1.4, or pCDNA3.1-Zeo(+) using jetPEI (Polyplus Transfection, Illkirch, France) following the manufacturer’s instructions. 48 hours after transfection, cells were fixed, stained, and imaged as described in the section **Confocal microscopy**.

### DETUBULATION

Isolated cardiomyocytes were detubulated by osmotic shock using formamide based on a previously described method^47^. Briefly, cardiomyocytes were resuspended in 2 mL 1.5 M formamide solution for 15 minutes and washed twice with extracellular solution (see “Electrophysiology” section).

### ELECTROPHYSIOLOGY

Whole-cell sodium currents (*I*_Na_) were recorded at 22-23°C using a VE-2 amplifier (Alembic Instruments Inc., Montréal, Canada). Borosilicate glass pipettes were pulled to a series resistance of ~2 MΩ. Recordings were visualized with pClamp software (version 8, Axon Instruments, Union City, California), and analyzed with pClamp and OriginPro (version 7.5, OriginLab Corp., Northampton, Massachusetts). Current densities (pA/pF) were calculated by dividing the peak current by the cell capacitance. Cells were bathed in extracellular solution containing (in mM) 5 NaCl, 125 NMDG-Cl, 5.6 CsCl, 5 BaCl2, 1 MgCl_2_, 10 HEPES, and 5 glucose (pH 7.4, CsOH adjusted). Internal solution contained (in mM) 130 KCl, 4 Mg-ATP, 12 NaCl, 1 MgCl_2_, 1 CaCl_2_, 10 EGTA, 10 HEPES (pH 7.2, KOH adjusted).

### STATISTICS

Data are presented as means ± standard deviation. Differences between two groups were analyzed in GraphPad Prism by two-tailed T-tests if the data were normally distributed, and Mann-Whitney tests if the data were not. *P* < 0.05 was considered statistically significant. No explicit power analysis was used. Three (SMLM) or five (electrophysiology) mice per genotype were used as biological replicates, providing sufficient statistical power. For SMLM, 47/24 (wild type), 26/27 (*mdx*), and 13/19 (ΔSIV) cells were imaged on intracellular and surface imaging planes, respectively, also providing sufficient statistical power. For individual experiments, cells from different genotypes were always isolated, or stained and imaged side by side. We performed 5/4 (wild type), 3/3 (*mdx*), and 2/3 (ΔSIV) technical replicates for intracellular and surface imaging, respectively. A technical replicate is defined as one round of staining and imaging. For electrophysiology, currents were recorded in seven control and seven detubulated cells from five wild-type mice. Each mouse counts as one technical replicate as data from each mouse were collected on one experiment day.

#### Exclusion and inclusion criteria

Regarding SMLM data, cells with irregular Bin1 staining were excluded, as they indicated a technical problem and/or T-tubular remodeling. All other cells were included as α-actinin and Na_v_1.5 stainings showed consistent results. No outliers were excluded.

Regarding electrophysiology data, we only included recordings with gigaseal and high-quality voltage clamp. Among those, no outliers were excluded.

## ACKNOWLEDGMENTS

The authors express their gratitude to Marta Pérez-Hernández Durán, Alejandra Leo-Macias, and Yandong Yin (NYUMC), and to Maria Essers and Sabrina Guichard (University of Bern) for their stellar technical assistance.

HA and SHV acknowledge funding from the Swiss National Science Foundation (grant no. 310030_165741 [HA] and P1BEP3_172237 [SHV]).

## AUTHOR CONTRIBUTIONS

SHV, HA, and JSR contributed conception of the study; SHV, JSR, EAP, ER, and MD contributed methodology; SHV and JSR performed experiments; SHV contributed validation, data curation, project administration, visualization, and writing–original draft; SHV and JSR performed formal analysis and investigation; EAP, ER, MD, and HA contributed resources; JSR, EAP, ER, MD, and HA supervised; SHV and HA contributed funding acquisition; and all authors contributed writing–review & editing.

## SOURCE DATA

**Figure 2d – Source data 1.** This spreadsheet contains the percentages of Na_v_1.5 clusters which edges are 50 nm from the edge of the nearest α-actinin cluster at the lateral membrane of wild-type cardiomyocytes, and the corresponding statistical analyses. Values from original images are compared to those from simulations in which Na_v_1.5 clusters are redistributed over the respective image either randomly or with high affinity for α-actinin.

**Figure 3d-f – Source data 2.** This spreadsheet contains data on Na_v_1.5 cell surface expression (pertaining to **Fig 3d**), frequency distribution of edge distances from any α-actinin cluster to the closest Na_v_1.5 cluster (pertaining to **Fig 3e**), and the percentage of Na_v_1.5 clusters within 50 nm from α-actinin at the lateral membranes of wild-type, *mdx*, and ΔSIV cells. Statistical analyses and descriptive statistics are given where applicable.

**Figure 4d-e – Source data 3.** This spreadsheet contains the values of the frequency distribution of edge distances from any Bin1 cluster to the closest Na_v_1.5 cluster (pertaining to **Fig 4d**), and the percentages of Na_v_1.5 clusters within 50 nm from Bin1 (pertaining to **Fig 4e**) in intracellular recordings of wild-type cardiomyocytes. Statistical analyses and descriptive statistics are also given.

**Figure 5 – Source data 4.** This spreadsheet contains values of cell capacitance, maximum sodium current, and sodium current density recorded in normal and detubulated wild-type cardiomyocytes. Statistical analyses are included.

**Figure 6 – Source data 5.** This spreadsheet contains values of Na_v_1.5 cluster density and frequency distributions of edge distances from any Bin1 cluster to the closest Na_v_1.5 cluster in intracellular planes in wild-type, *mdx*, and ΔSIV cells. Statistical analyses and descriptive statistics are included.

**Supplemental figure 2 – Source file 6.** This spreadsheet contains values of cell size, average Na_v_1.5 cluster size, Na_v_1.5 cluster solidity and circularity of wild-type, *mdx*, and ΔSIV cardiomyocytes at surface (pertaining to **Supplemental Fig 2a**) and intracellular imaging planes (pertaining to **Supplemental Fig 2b**). Statistical analyses are included.

**Supplemental figure 3 – Source file 7.** This spreadsheet contains values of Na_v_1.5 clusters within 50 nm from α-actinin at the cell surface (pertaining to **Supplemental Fig 3a**) and Bin1 inside the cell (pertaining to **Supplemental Fig 3b**). Values are compared between original images and simulations in which Na_v_1.5 clusters are redistributed either randomly or with high affinity for α-actinin or Bin1, respectively. Statistical analyses are also given.

## SUPPLEMENTAL FIGURES

**Supplemental figure 1.**
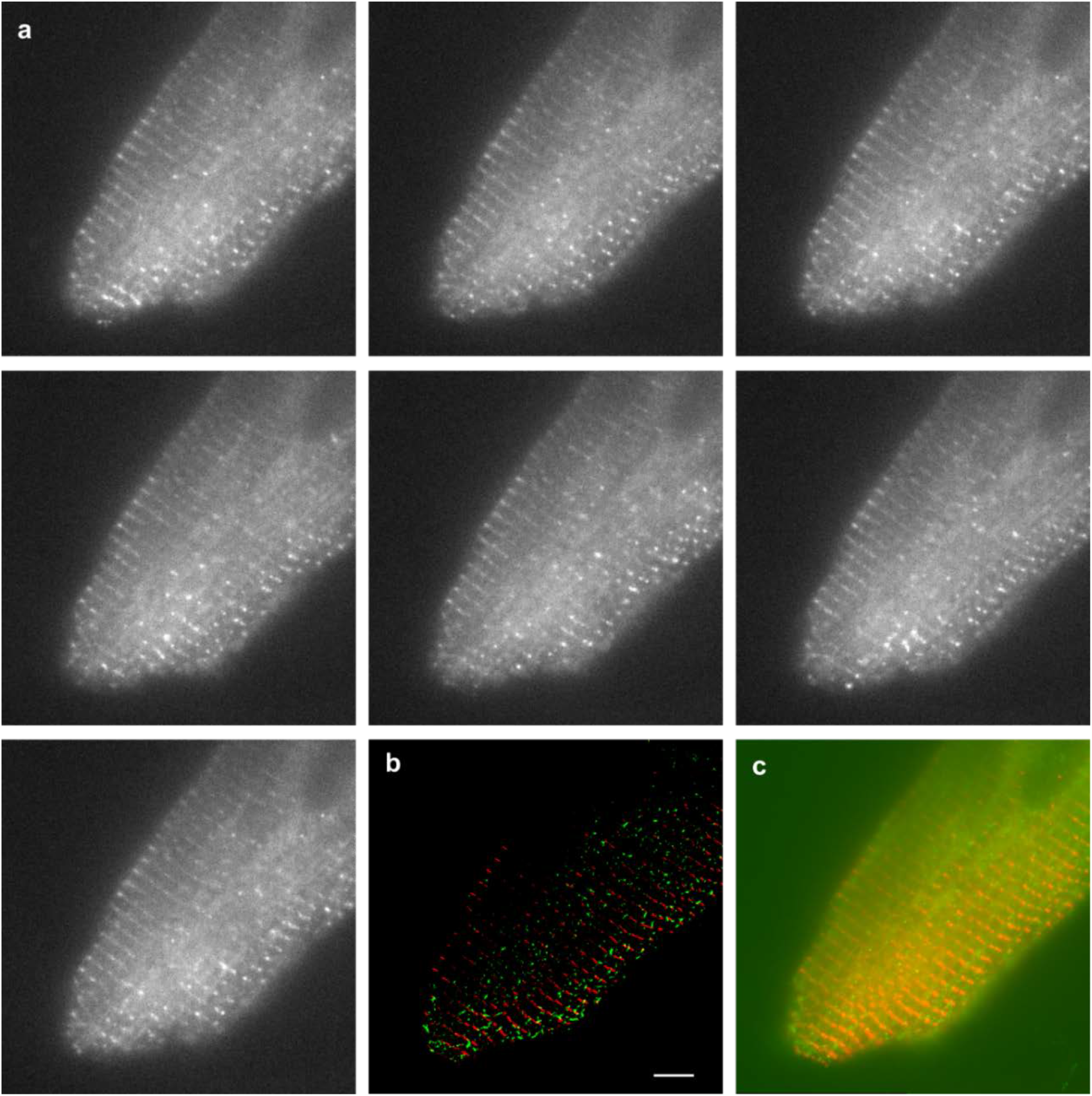
Fluorophores blink during SMLM image acquisition. **a**, Frames from a recording of a cardiomyocyte stained with anti-α-actinin antibodies directly conjugated to AlexaFluor 647 fluorophores and excited with a 639 nm laser. The fluorophores release photons intermittently – “blink” – due to the oxygen scavenging imaging buffer (see Methods). **b**, Reconstructed image depicting α-actinin in red and Na_v_1.5 in green. Note that the fluorophores on the left side of the cell that did not blink, characterized by a faint striated pattern in all panels of (**a**), are not reconstructed in the final SMLM image (**b**). Scale bar 5 μm.

**Supplemental figure 2.**
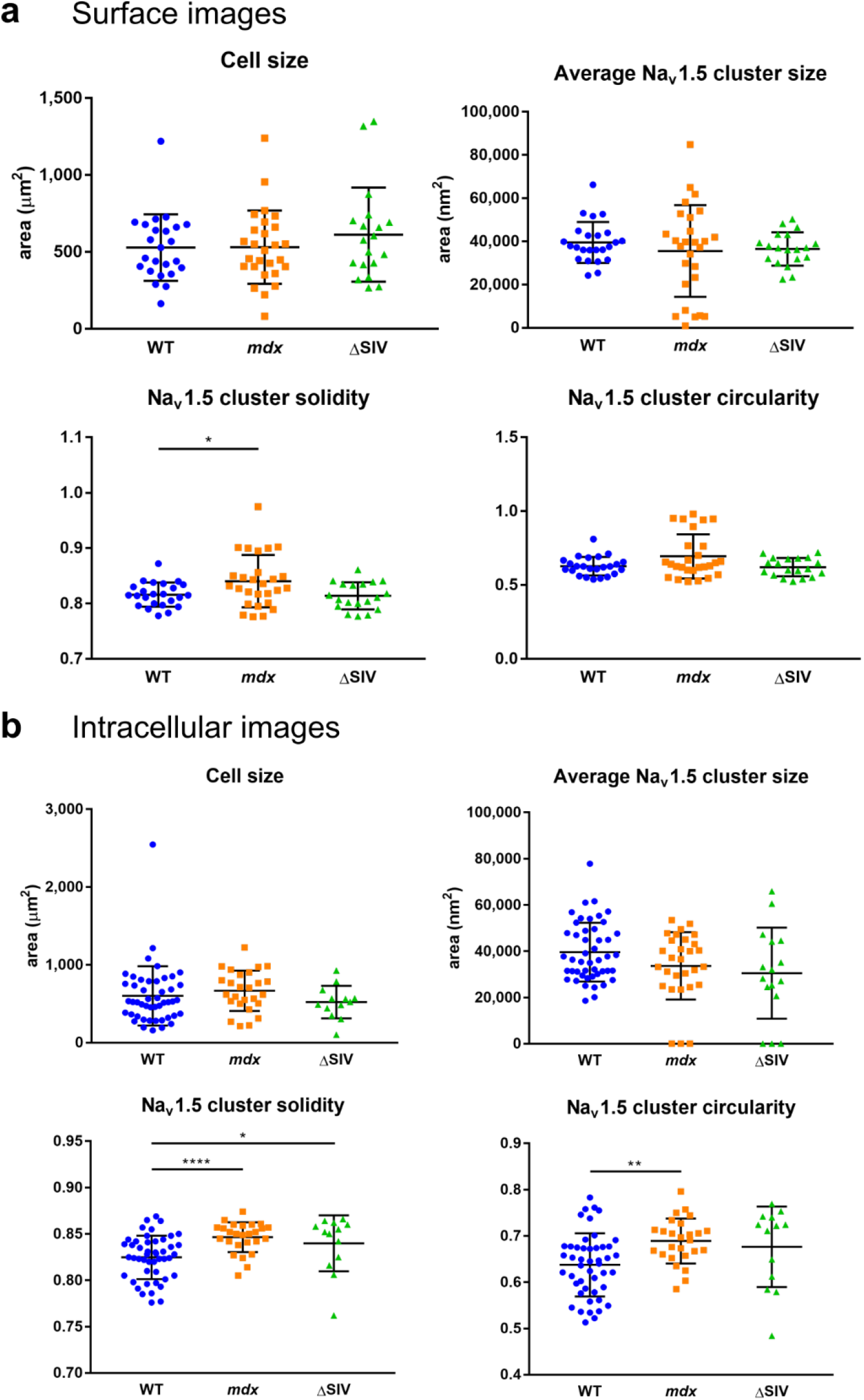
Cell and Na_v_1.5 cluster properties of SMLM images. For all images from wild-type, *mdx*, and ΔSIV cells at the lateral membrane (**a**) and inside the cell (**b**), this figure shows cell size, and cluster size, circularity, and solidity of Na_v_1.5 clusters per cell. Perfectly circular and solid clusters give value 1. Solidity indicates the ratio of the particle area to the area of the convex hull of the particle. **a**, Wild type *N* = 3, *n* = 24; *mdx N* = 3, *n* = 27; ΔSIV *N* = 3, *n* = 19. **b**, Wild type *N* = 3, *n* = 39; *mdx N* = 3, *n* = 20; ΔSIV *N* = 3, *n* = 11. *, *p* = 0.041 (**a**, solidity); *p* = 0.021 (**b**, solidity); **, *p* = 0.001; ****, *p* < 0.0001, Mann Whitney test.

**Supplemental figure 3.**
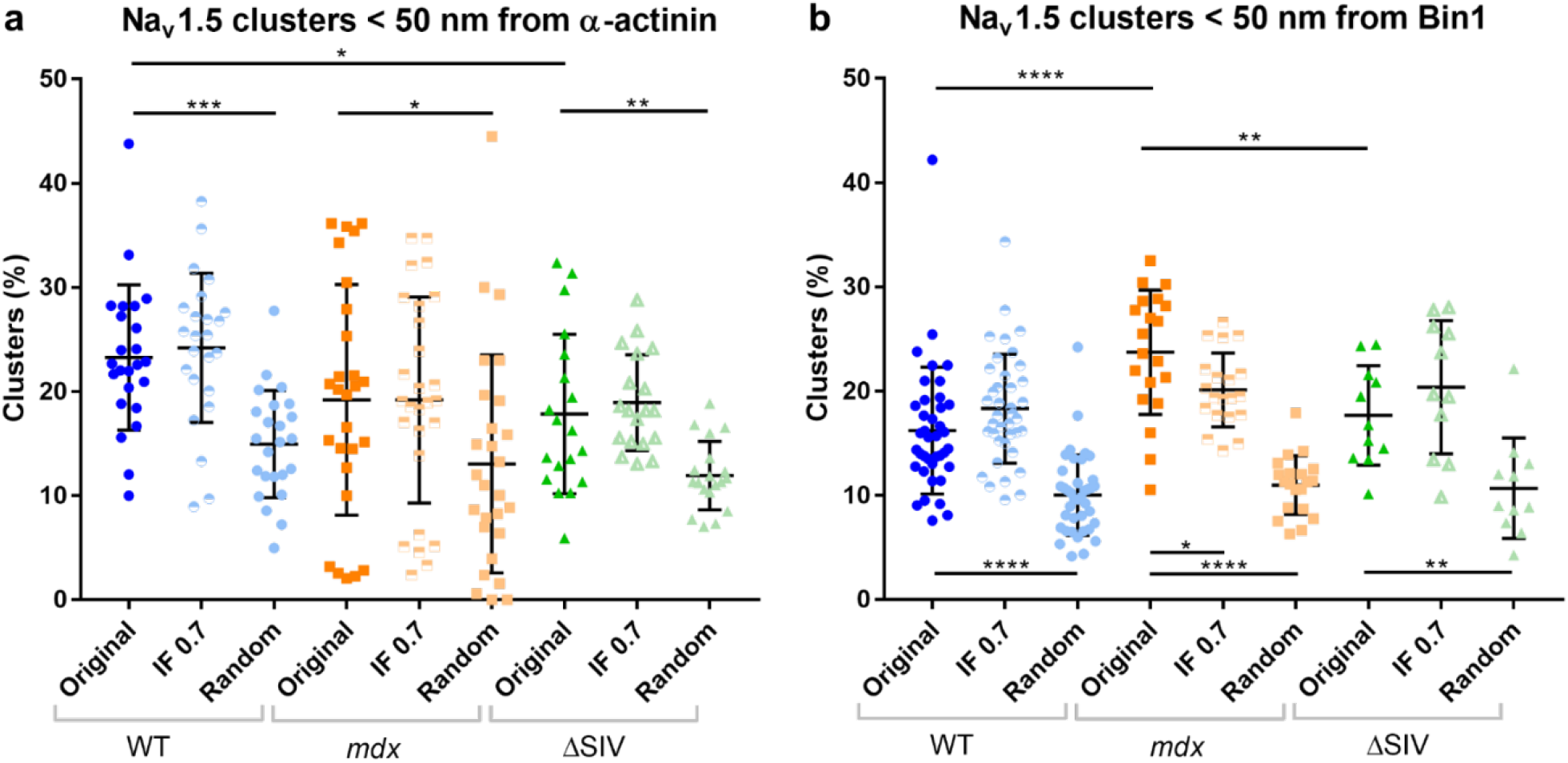
Comparing cluster distances of experimental SMLM images to simulations. Proportion of Na_v_1.5 clusters within 50 nm from α-actinin at the lateral membrane (**a**) and Bin1 inside the cell (**b**). SMLM images from wild-type (blue), *mdx* (orange), and ΔSIV (green) cells are compared to their respective high affinity (IF 0.7) and random simulations. **a**, Wild type *N* = 3, *n* = 24; *mdx N* = 3, *n* = 27; ΔSIV *N* = 3, *n* = 19. **b**, Wild type *N* = 3, *n* = 39; *mdx N* = 3, *n* = 20; ΔSIV *N* = 3, *n* = 11. *, *p* = 0.0194 (**a**, ΔSIV-WT), *p* = 0.017 (**b**, *mdx*); **, *p* = 0.0074 (**a**, ΔSIV) *p* = 0.0019 (**b**, ΔSIV), *p* = 0.0095 (**b**, *mdx*-ΔSIV); ***, *p* < 0.001, ****, *p* < 0.0001, unpaired T-test (**b**, wild type) and Mann Whitney test (all others).

**Supplemental figure 4.**
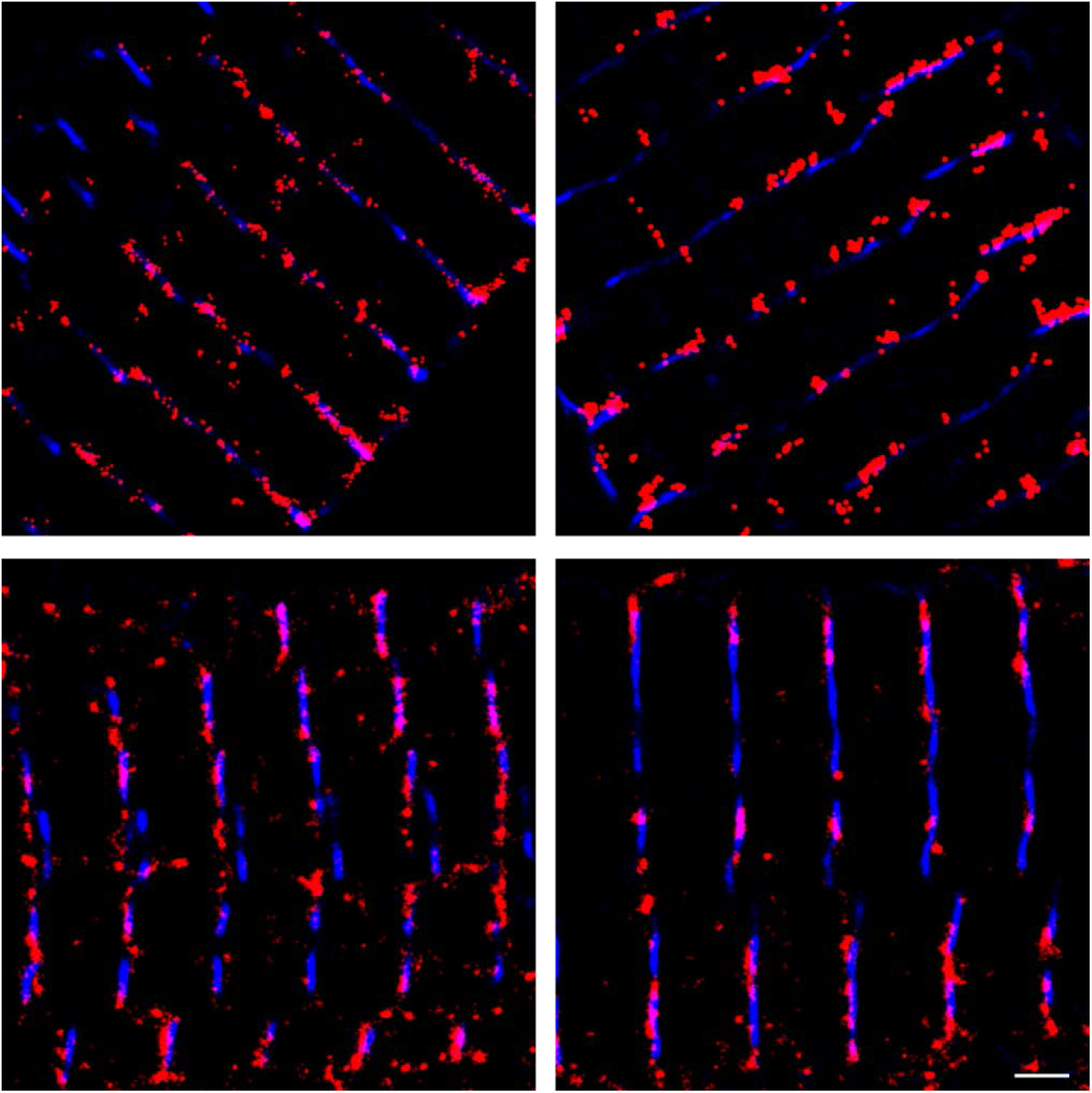
Bin1 antibody validation. Representative SMLM images of Bin1 (red) and α-actinin (blue) showing close proximity of Bin1 to α-actinin. This indicates that the anti-Bin1 antibody is most likely specific, as Bin1 binds the T-tubular membrane, and most T-tubules run along α-actinin lines, which constitute the sarcomeric Z-discs. Bin1 signal in between α-actinin lines may indicate axial T-tubular branches. *N* = 3, *n* = 57; scale bar 1 μm.

**Supplemental figure 5.**
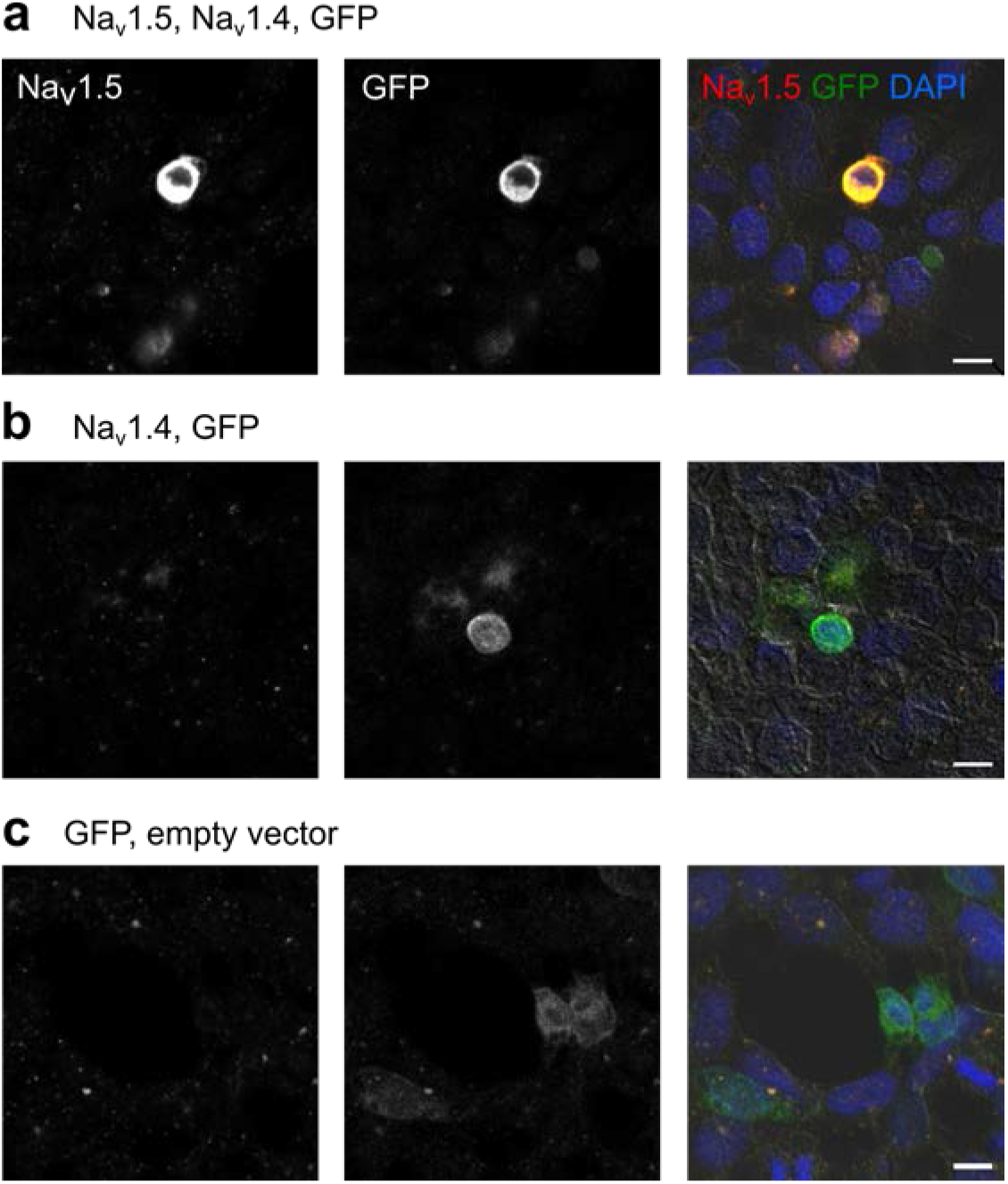
Anti-Na_v_1.5 antibody (Pineda) does not recognize Na_v_1.4. Representative confocal images of TsA-201 cells transfected with (**a**) Na_v_1.5, Na_v_1.4, and GFP; (**b**) Na_v_1.4 and GFP; (**c**) and GFP and empty vector. Cells are stained for Na_v_1.5 (Pineda antibody; red), GFP (green), and nuclei (DAPI; blue). Scale bar 10 μm.

